# EfpA is required for re-growth of *Mycobacterium tuberculosis* following isoniazid exposure

**DOI:** 10.1101/2023.08.31.555697

**Authors:** Adam H. Roberts, Christopher W. Moon, Valwynne Faulkner, Sharon L. Kendall, Simon Waddell, Joanna Bacon

**Affiliations:** Discovery Group, Research and Evaluation, UKHSA, Porton Down, Salisbury, SP4 0JG; Centre for Emerging, Endemic and Exotic Diseases, Pathobiology and Population Sciences, Royal Veterinary College, AL9 7TA; Global Health and Infection, Brighton and Sussex Medical School, University of Sussex, BN1 9PX, UK

**Keywords:** EfpA, efflux pumps, *Mycobacterium tuberculosis*, isoniazid, re-growth, relapse, NADH/NAD^+^

## Abstract

Efflux of antibiotics is a survival strategy in bacteria. *Mycobacterium tuberculosis* has approximately sixty efflux pumps, but little is known about the role of each pump, or which moieties that they efflux. The putative efflux pump, EfpA, is a member of the major facilitator superfamily that has been shown to be essential by saturation transposon mutagenesis studies. It has been implicated in the efflux of the frontline drug isoniazid (INH) in *M. tuberculosis*. This is supported by evidence from transcriptional profiling that *efpA* is induced in response to INH exposure. However, its role in physiology and adaptation of *M. tuberculosis* to antibiotics, have yet to be determined. Here, we describe the repression of *efpA* using CRISPR interference and the direct effect of this on the ability of *M. tuberculosis* to survive exposure to INH over a 45-day time-course. We determined that wild-type levels of *efpA* were required for the recovery of *M. tuberculosis* cultures following INH exposure and that, after 45-days of INH exposure, no viable colonies were recoverable from *efpA*-repressed *M. tuberculosis* cultures. We postulate that EfpA is required for the recovery of *M. tuberculosis* following INH-exposure and that EfpA may have a role in the development of resistance, during treatment and contributes to relapse in patients.

## Introduction

Infection with *Mycobacterium tuberculosis* once again is the leading cause of death from a single infectious agent. In 2021, approximately 10.6 million people became ill with tuberculosis (TB) and 1.6 million people died from TB world-wide (WHO, 2022). The worsening of multi-drug resistance in *M. tuberculosis*, defined as resistance to at least isoniazid (INH) and rifampicin, has also compounded the burden of TB infections globally with 450,000 cases estimated to have occurred in 2021 alone (WHO, 2022). The lengthy and toxic nature of TB multi-drug therapy highlights a need for more regimens with lower toxicity and shortened treatment times. This requires greater insight into new and existing mycobacterial drug targets for which promising new therapies can emerge to combat the global TB burden.

Although drug resistance in *M. tuberculosis* is predominately caused by alterations in drug activating enzymes or drug targets through DNA mutation, there is increasing evidence of enhanced expression of efflux pumps (including EfpA) in multidrug-resistant clinical isolates (Li *et al*., 2015; Kardan-Yamchi *et al*., 2019; Shahi *et al*., 2021) that may contribute to the development of drug resistance. Studies by Machado *et al*. (2012) and Rodrigues *et al*. (2012) demonstrated a relationship between the overexpression of genes encoding efflux pumps and increased efflux activity, with an increased frequency in *katG* mutations. This supports the hypothesis that efflux of an antibiotic provides a window of opportunity for genetically encoded drug-resistance to emerge. Therefore the discovery of molecules that inhibit the action of pumps, potentially extending the life of current and new therapies, would be an advantage in combating both drug-susceptible and drug-resistant *M. tuberculosis* infections (Johnson *et al*., 2019; Laws, Jin, & Rahman, 2022).

The efflux pump, EfpA, which belongs to the major facilitator family, is an essential pump in *M. tuberculosis* as demonstrated by transposon mutagenesis studies (Sassetti, Boyd, & Rubin, 2003; Griffin *et al*., 2011; Dejesus *et al*., 2017; Minato *et al*., 2019). This essentiality brings challenges in that the function cannot be studied by deleting the gene. The first study to identify and characterise the gene, *efpA*, was by Doran *et al*., in 1997, who found that *efpA* encodes a putative efflux protein that is predicted to be highly related to the QacA transporter family, with orthologues in *M. marinum, M. leprae, M. smegmatis*, and *M. bovis*. It is possible that the regulation of *efpA* has a role in the survival of *M. tuberculosis* in the acidic compartment of the macrophage as its expression is reduced under acidic conditions (Fisher, Plikaytis, & Shinnick, 2002). Whole genome microarray studies also revealed that in *M. tuberculosis, efpA* is more highly expressed during INH exposure *in vitro* (Wilson *et al*., 1999; Boshoff *et al*., 2004; Waddell *et al*., 2004; Gupta *et al*., 2010; Jeeves *et al*., 2015), and in sputa (Honeyborne *et al*., 2016).

Populations of *M. tuberculosis* exposed to INH, *in vitro*, exhibit a substantial loss in bacterial viability during the first two days followed by a period of re-growth (Gumbo *et al*., 2007; de Steenwinkel *et al*., 2010; Jeeves *et al*., 2015; Hendon-Dunn *et al*., 2019), which is not entirely explained by an increase in the INH-resistant mutant frequency (Jeeves *et al*., 2015). This biphasic response has also been observed in *M. tuberculosis*-infected guinea pigs treated with INH and was found to be associated with the emergence of antibiotic-tolerant persisting populations that were not INH-resistant mutants (Ahmad *et al*., 2009). Efflux has been suggested as a mechanism that might contribute to *M. tuberculosis* drug tolerance and/or INH resistance (Schmalstieg *et al*., 2012).

There is direct evidence for the role of *efpA* in the response of *M. tuberculosis* to the standard regimen during the early stages of treatment; *efpA* was induced in TB patient sputa 3 days after the start of drug therapy (Honeyborne *et al*., 2016). It would be informative for therapeutic targeting of efflux mechanisms in *M. tuberculosis* if we could understand the contributions of efflux to drug tolerance and emergence of genotypic resistance, and more specifically, the role of the efflux pump, EfpA, in mediating efficacy of the frontline drug INH. Since *efpA* is an essential gene, we used CRISPRi knock-down to determine the significance of EfpA in the viability of *M. tuberculosis* when exposed to INH, over time, *in vitro*.

## Materials and methods

### Bacterial strains, plasmids, and culture conditions

*M. tuberculosis* H37Rv strains were grown in Difco Middlebrook 7H9 medium (Sigma-Aldrich), supplemented with 0.5% glycerol (VWR), 0.2% tween-80 (Sigma-Aldrich), and 10% oleic-acid dextrose catalase (OADC) growth enrichment supplement (UKHSA-Porton Down Media services). 7H10 Middlebrook medium was utilised for the growth of *M. tuberculosis* on agar, which was supplemented with 0.5% glycerol and 10% OADC growth enrichment supplement. Each CRISPRi culture contained the selection antibiotics kanamycin and hygromycin at final concentrations of 25 μg mL^-1^ and 50 μg mL^-1^, respectively. All liquid cultures were grown at 37°C in a 200 RPM shaking incubator. Each culture was grown from frozen stocks for 10 days prior to inoculation and standardised to an OD_540nm_ of 0.05 for growth curves and time-kill survival assays, and OD_540nm_ of 0.1 for cultures where RNA was extracted for qRT-PCR. *M. tuberculosis* on Middlebrook 7H10 solid medium was incubated statically at 37°C for 3 weeks.

### Antibiotic stock production

Isoniazid (Sigma-Aldrich) and anhydrotetracycline (ATc) (Sigma-Aldrich) solutions were prepared by dissolving them in dimethyl sulfoxide (DMSO), whereas kanamycin (Sigma-Aldrich) and hygromycin (Sigma-Aldrich) solutions were prepared by dissolving the compounds in water. All drug stocks were filter sterilised using a 0.2 μm filter (Sartorius Stedim).

### Plasmid and strain construction

The extrachromosomal pRH2521 plasmid, encoding for single-guide RNA (sgRNA) scaffold sequence and the integrative pRH2502 plasmid, encoding for a deactivated Cas9 (dCas9) from *Streptococcus pyogenes* (Table 1), (both under the control of Tet-regulatory promoters) were used for *efpA* silencing. The two-plasmid CRISPRi system was developed for *M. tuberculosis* by Singh *et al*. (2016); plasmids pRH2521 and pRH2502 were gifts from Robert Husson (Addgene plasmid # 84380 and Addgene plasmid # 84379). The *efpA*-targeting sequence was cloned into the sgRNA using the method outlined by Larson *et al*. (2013), which was also performed by Faulkner *et al*. (2021) and Gibson *et al*. (2021). Initially, a 20nt DNA sequence was identified as a target for an sgRNA base-pairing sequence 98nt downstream of the start of the annotated *efpA* open reading frame (ORF) in *M. tuberculosis*, which was downstream of a protospacer adjacent motif (PAM) sequence ‘NGG’. For this sequence, a check was performed for complementarity relative to other genes in the *M. tuberculosis* genome using the nucleotide basic local alignment search tool (BLASTn) (Altschul *et al*., 1990). The 20nt (without PAM sequence) was reverse complemented to form the sgRNA sequence and combined with the dCas9 handle and terminator sequences *in silico* and the secondary structure was predicted using the M-fold tool (Zuker, 2003) to ensure that the dCas9 handle and terminator hairpins were likely to form correctly. Forward and reverse primers containing the 20nt sequence with ‘overhang’ sequences included at the 5’ end, forward primers with “GGGA” sequence and reverse primers with “AAAC” to allow for ligation into the pRH2521 plasmid during cloning. These oligonucleotides were annealed together and the insert was cloned into pRH2521 plasmid using the BbsI restriction enzyme (NEB) and *E. coli* DH5α, as per methodology outlined by Singh *et al*. (2016). 1 μg of the *efpA-*targeting sgRNA pRH2521 plasmid was electroporated into an electrocompetent *M. tuberculosis* H37Rv_dCas9_ strain (containing pRH2502 plasmid), following a published protocol (Parish & Brown, 2009) using a pulse of 2.5kV, 25μF with resistance set at 1000Ω using a Gene Pulser Xcell (Bio-Rad). The successful transformants were selected for using 7H10 medium containing 25 μg mL^-1^ kanamycin and 50 μg/mL hygromycin. The *M. tuberculosis* H37Rv strain containing pRH2521, encoding for the *efpA*-targeting sgRNA, is referred to as the *<u>efpA</u>* knockdown strain (*efpA*^KD^). Control *M. tuberculosis* H37Rv strains contained a dCas9-encoding pRH2502 plasmid only for electroporation or a strain which contained pRH2502 in addition to a pRH2521 plasmid that encoded an sgRNA with a random base-pairing sequence that was not specific to any *M. tuberculosis* gene target (the latter is referred to as “control strain”).

**Table 1:**
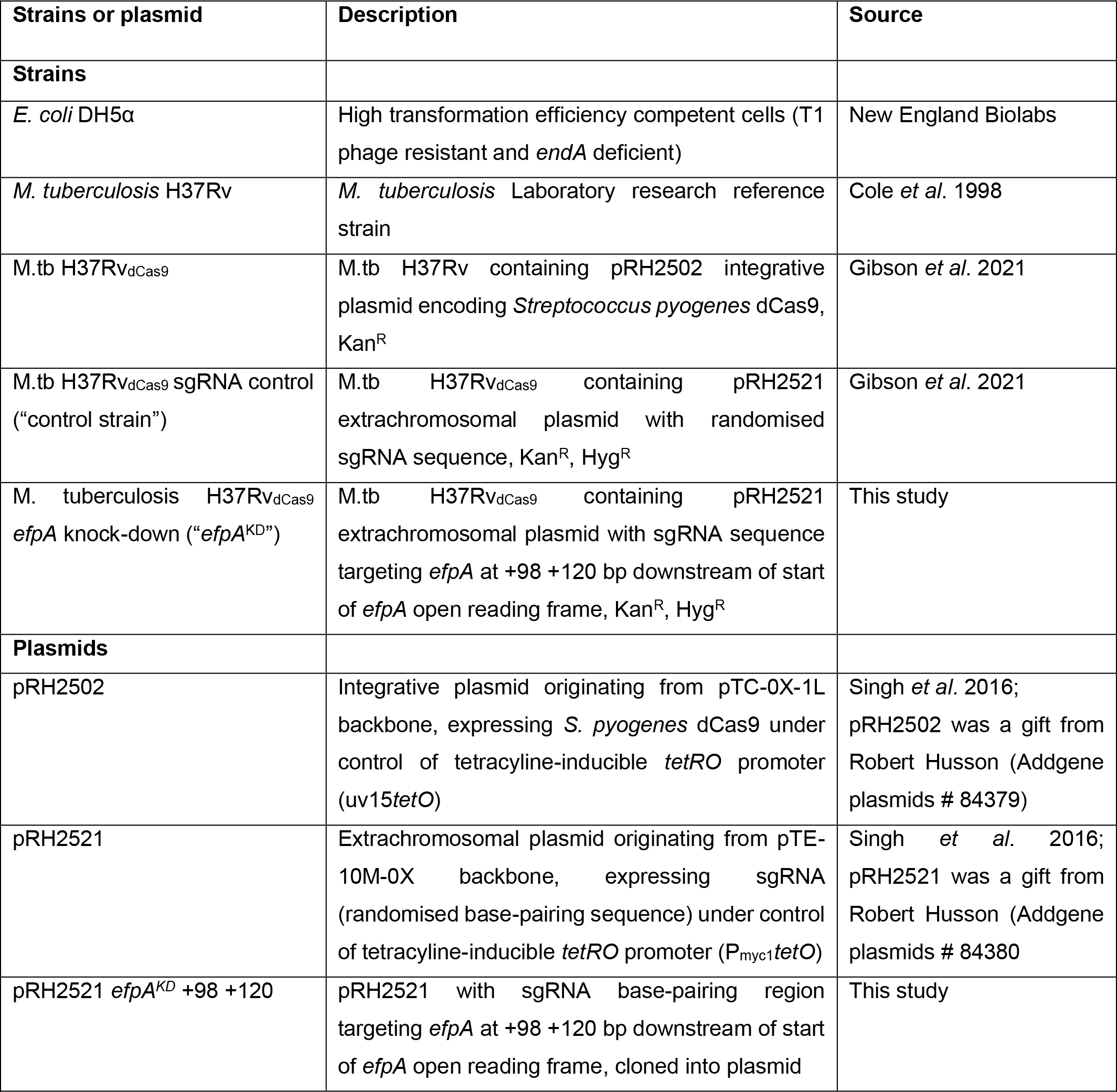
Strains and plasmids used.

| Strains or plasmid | Description | Source |
| --- | --- | --- |
| **Strains** |  |  |
| *E. coli* DH5α | High transformation efficiency competent cells (T1 phage resistant and *endA* deficient) | New England Biolabs |
| *M. tuberculosis* H37Rv | *M. tuberculosis* Laboratory research reference strain | Cole *et al.* 1998 |
| M.tb H37Rv<sub>dCas9</sub> | M.tb H37Rv containing pRH2502 integrative plasmid encoding *Streptococcus pyogenes* dCas9, Kan<sup>R</sup> | Gibson *et al.* 2021 |
| M.tb H37Rv<sub>dCas9</sub> sgRNA control (“control strain”) | M.tb H37Rv<sub>dCas9</sub> containing pRH2521 extrachromosomal plasmid with randomised sgRNA sequence, Kan<sup>R</sup>, Hyg<sup>R</sup> | Gibson *et al.* 2021 |
| *M. tuberculosis* H37Rv<sub>dCas9</sub> *efpA* knock-down (“*efpA*<sup>KD</sup>”) | M.tb H37Rv<sub>dCas9</sub> containing pRH2521 extrachromosomal plasmid with sgRNA sequence targeting *efpA* at +98 +120 bp downstream of start of *efpA* open reading frame, Kan<sup>R</sup>, Hyg<sup>R</sup> | This study |
| **Plasmids** |  |  |
| pRH2502 | Integrative plasmid originating from pTC-0X-1L backbone, expressing *S. pyogenes* dCas9 under control of tetracycline-inducible *tetRO* promoter (uv15*tetO*) | Singh *et al.* 2016; pRH2502 was a gift from Robert Husson (Addgene plasmids # 84379) |
| pRH2521 | Extrachromosomal plasmid originating from pTE-10M-0X backbone, expressing sgRNA (randomised base-pairing sequence) under control of tetracycline-inducible *tetRO* promoter (P<sub>myc1</sub>*tetO*) | Singh *et al.* 2016; pRH2521 was a gift from Robert Husson (Addgene plasmids # 84380) |
| pRH2521 *efpA*<sup>KD</sup> +98 +120 | pRH2521 with sgRNA base-pairing region targeting *efpA* at +98 +120 bp downstream of start of *efpA* open reading frame, cloned into plasmid | This study |

### ATc-induction of *efpA* repression

The *efpA*^KD^ and control strain were grown in Middlebrook 7H9 medium over 14-days in a volume of 13 mL, in universal tubes, across three biological replicates. At the point of inoculation, a final concentration of ATc at 200 ng mL^-1^ was added to the cultures. At intervals, culture was collected, spun at 6,000 RPM for 10 minutes and the pellet was resuspended in phosphate buffered saline (PBS) pH 7.4. From the resuspended sample, viable counts (CFU mL^-1^) were determined by serially diluting culture in PBS and plating out onto 7H10 agar using the Miles-Misra technique. The plates were incubated for 3 weeks at 37°C in a static incubator prior to enumeration of the viable count.

### *M. tuberculosis* cultures exposed to isoniazid

To assess the impact of INH exposure, the *efpA*^KD^ and control strains were grown in Middlebrook 7H9 medium over 45-days in three replicate cultures. At the point of inoculation, ATc (final concentration of 200 μg mL^-1^) and/or isoniazid (final concentration of 0.5 μg mL^-1^) were added to the cultures. The time-kill survival experiments also included a no-drug control condition. At intervals, culture was collected, spun at 6,000 RPM for 10 minutes, and the pellet was resuspended in PBS. Viability was measured using total viable counts (CFU mL^-1^) as described above.

### RNA extraction and purification

RNA was extracted from 10 mL of culture sampled at intervals of 0, 1, 3 and 7-days post ATc-addition. 40 mL of guanidine thiocyanate (GTC) lysis solution (comprised of 5M GTC, 0.5% lauryl sarcosine, 25mM tri-sodium citrate, 0.5% tween-80, and 0.05% 2-mercaptoethanol) was added to the culture samples and incubated for 1 hour at room temperature. The cultures were spun at 3,000 RPM for 10 minutes, the supernatant was removed, and the *M. tuberculosis* pellet was resuspended in 1.2 mL Trizol solution (Thermo) and transferred to lysing matrix B beads (0.5 mL of 0.1mm) (MP bio) and placed into a mini bead-beater (Biospec) and the cells were disrupted at full power for 50 seconds. The solution was spun at 13,000 RPM for 10 minutes and resuspended in 240 μL 24:1 chloroform: isoamyl alcohol (Sigma-Aldrich) and shaken vigorously for 20 seconds. Following centrifugation at 13,000 RPM for 10 minutes, the aqueous phase was resuspended in 600 μL chloroform: isoamyl alcohol, centrifugated again and the step was repeated. The aqueous phase was added to 600 μL of isopropanol (Sigma-Aldrich) with 60 μL sodium acetate (Sigma-Aldrich) and incubated overnight at -70°C to precipitate the nucleic acids. The sample was washed with ethanol, resuspended in nuclease-free water and purified using a Qiagen RNeasy kit with additional on-column DNase (Qiagen) digestion. RNA quantity and purity were assessed using a spectrophotometer (Nanodrop; Thermo).

### Quantitative reverse transcription-PCR (RT-qPCR)

The absolute quantification method was used to determine the number of transcripts (copy number) of *efpA* in each sample, relative to an endogenous housekeeping control gene, *sigA*. To achieve a copy number standard curve for absolute quantification, full gene fragments of *efpA* and *sigA* amplified from *M. tuberculosis* H37Rv genomic DNA using the PCR primers outlined in Table 2, GoTaq green polymerase (Promega), and DMSO (Sigma-Aldrich), using conditions of 98°C for 2 minutes, followed by 35 cycles of 94°C for 30 seconds, 59°C for 30 seconds and 72°C for 2 minutes, with a final cycle of 72°C for 10 minutes. The PCR product was purified using a PCR clean-up kit (Qiagen) and the DNA concentration was quantified using a Qubit broad range quantification assay (Thermo). From the quantity and gene length, the copy number of *efpA* and *sigA* was calculated using the Thermo Fisher copy number calculator. The copy number values were standardised to 10^7^ and used to generate a standard curve of *efpA* and *sigA* abundance. The reverse transcription reactions to synthesise cDNA consisted of 500 ng total RNA, 1 μL Superscript III (Thermo), 4 μL First Strand Synthesis Buffer (Thermo), 1 μL 10mM dNTP mixture (Thermo), and 1 μL random primers (Thermo). The reactions also featured a no reverse transcriptase control for each sample. For the qRT-PCR reactions, 10 μL Taqman™ universal master-mix II with UNG (Thermo), 5 μL of cDNA/control DNA fragment, 1 μL forward and reverse primers, 0.5 μL Taqman probe, and nuclease-free water to reach a total reaction volume of 20 μL in a 96-well MicroAmp plate (Thermo). The reaction cycling conditions used were 2 minutes at 95°C followed by 45 cycles of 95°C for 30 seconds and 60°C for 30 seconds using the Quantstudio™ 7 Real-Time PCR system (Thermo). Each qRT-PCR reaction run featured non-template controls. The data was analysed using the QuantStudio™ Real-Time PCR Software (Thermo) and *efpA* copy numbers were normalised as a ratio to the *sigA* copy number. The values from the ATc-induced *efpA*^KD^ were analysed relative to the non-induced *efpA*^KD^ and the ATc-induced control strain.

**Table 2:**
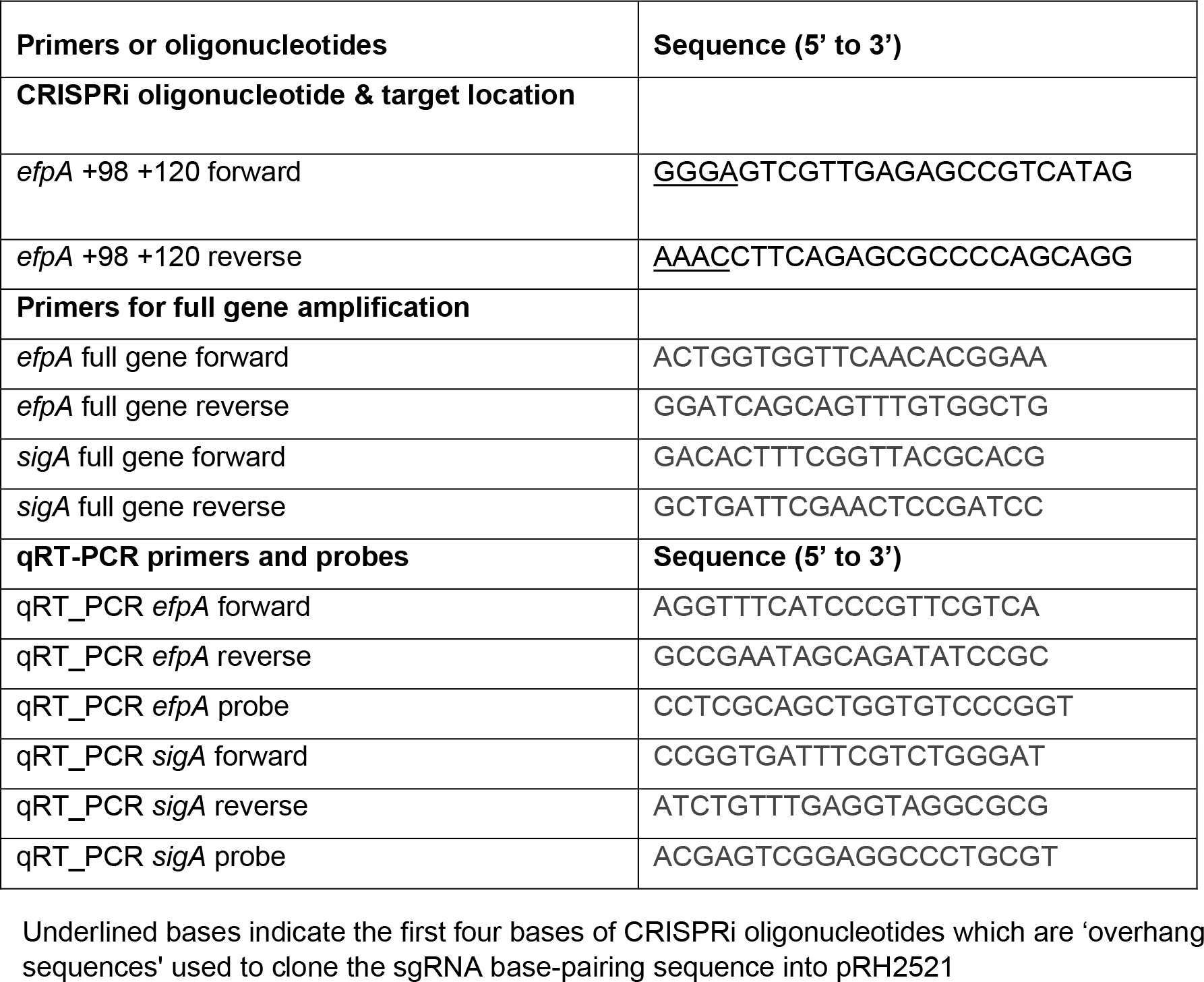
Primers and oligonucleotides used in this study.

### Scanning electron microscopy

*M. tuberculosis* CRISPRi cultures were fixed at day 7 post ATc-induction using a final concentration of 4% formaldehyde, centrifugated at 6,000 RPM for 10 minutes and resuspended in water. Formaldehyde-fixed cells were sent to the Scanning Electron Microscopy (SEM) department at UKHSA-Porton Down for imaging. The imaging process involved the immobilisation of cells onto a glass cover slip where cells were further fixed using osmium tetroxide for 2 hours at room temperature followed by dehydration using a graded series ethanol-exposure. Samples were then treated with hexamethyldisilane and air-dried and finally the conductive coating of 20 nm gold was applied using a fine grain ion-beam sputter coater, prior to imaging using a Zeiss Sigma 300VP Scanning Electron Microscope.

### Statistical methods

For the growth curve (Fig. 2), a two-way ANOVA was applied to identify any significant differences in the total viable count (CFU mL^-1^) across the time-course between strains with or without ATc-induction. For the isoniazid time-kill experiments (Fig. 5), a three-way ANOVA was utilised to compare viability (CFU mL^-1^) between strains and between ATc-induction and isoniazid-exposure conditions. All viable counts were log_10_ transformed prior to statistical analyses. ANOVAs were performed using RStudio software (version 1.3.1056).

## Results

### Targeted repression of *efpA* results in a loss of viability in *M. tuberculosis*

The aim of this study was to determine the role of EfpA in the response of *M. tuberculosis* to INH, *in vitro*. Since *efpA* is an essential gene in *M. tuberculosis*, an “*efpA* knock-down strain” (*efpA*^KD^) was generated by CRISPRi and induced with anhydrotetracycline (ATc), using the two plasmid-dCas9 system developed by Singh *et al*. (2016). The sgRNA targeted *efpA* in a 20bp region between +98bp to +120bp relative to the start codon of the *efpA* coding region, downstream of a 3bp protospacer adjacent motif (PAM) required for dCas9 binding (Fig. 1). A “control strain” was generated that contained the same vectors but with a random sgRNA sequence, rather than an *efpA*-targeting sgRNA, as a comparator in all experiments (Singh *et al*., 2016).

**Figure 1:**
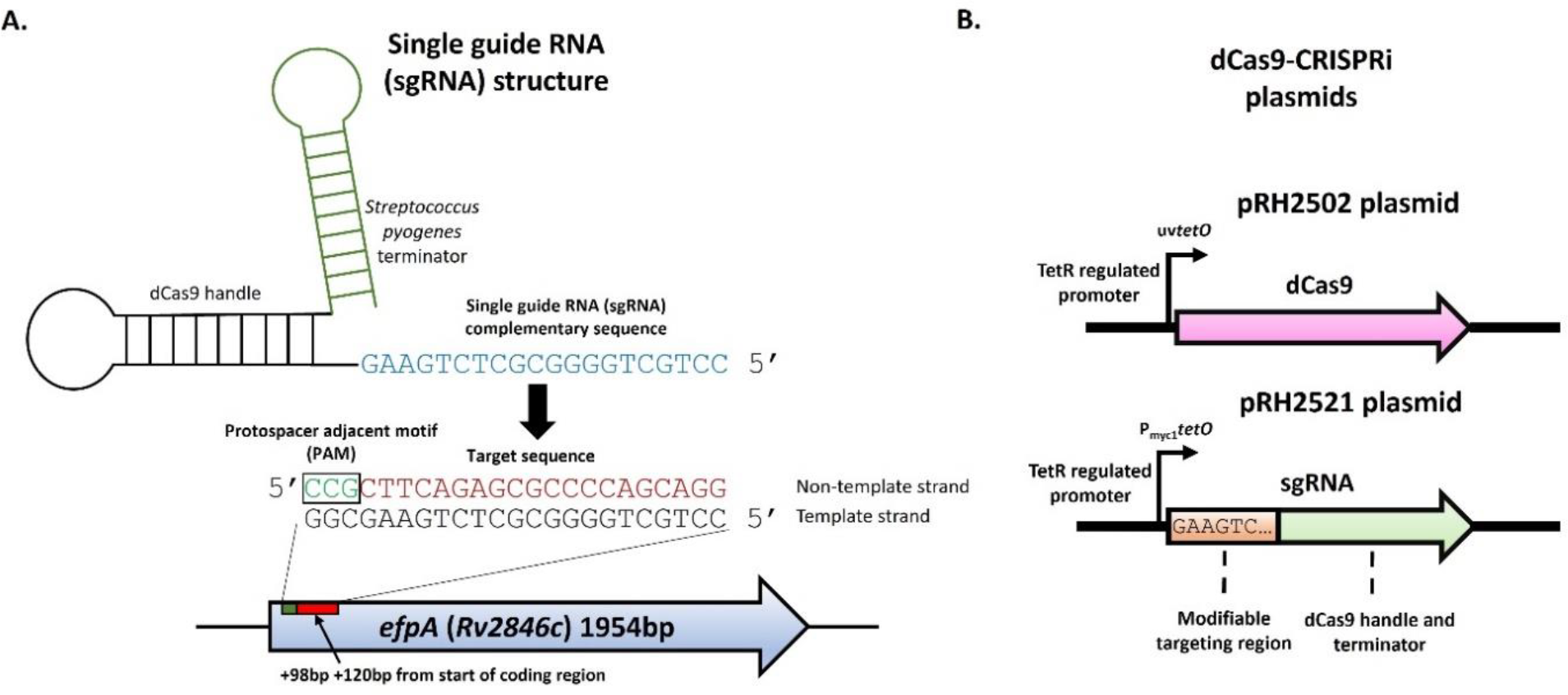
A two-plasmid dCas9-sgRNA CRISPRi system adapted to repress *efpA* expression. Schematic of the single-guide RNA chimeric structure consisting of the terminator region originating from *Streptococcus pyogenes*, the dCas9 handle region (to enable binding to dCas9 enzyme), and the sequence complimentary to the *efpA* target sequence at the 5’ end of the gene. The protospacer adjacent motif (PAM) sequence (CCN) is adjacent to the 5’ end of the target sequence (Panel A). Schematic of the CRISPRi genes on two plasmids coding for the CRISPRi system; pRH2502, which encodes the dCas9 endonuclease enzyme under the control of a uv*tetO* promoter, and pRH2521-*efpA*, which encodes the *efpA* sgRNA under the control of a p*myc1tetO* promoter (Panel B).

The viability of the *efpA*^KD^ was determined after ATc-induction compared to a non-induced state, in triplicate flask cultures, over a 14-day time-course in 7H9 Middlebrook medium. ATc was added to the respective cultures on day 0 of the time-course at a final concentration of 200 ng mL^-1^, which had been optimised as the lowest ATc concentration optimised to maximally reduce *efpA* expression. Samples were removed at days 0, 1, 2, 3, 4, 7, 9, 11 and 14, for total viable counts (Fig. 2), qRT-PCR (Fig. 3), and scanning electron microscopy (Fig. 4). The conditions, such as concentration of ATc and the time of addition, were optimised prior to these experiments.

The knock-down of *efpA*, resulted in a steep reduction in viability between days 2 and 4 of the time-course (Fig. 2) and there was a statistically significant difference between this response and the non-induced *efpA* knock down strain (*P* = <0.001). Despite a gradual increase in viability from day 7, the induced *efpA*^KD^ was unable to recover to the same level of viability as that of the non-induced strain at day 14. As expected, ATc-addition did not affect the viability of the control strain.

**Figure 2:**
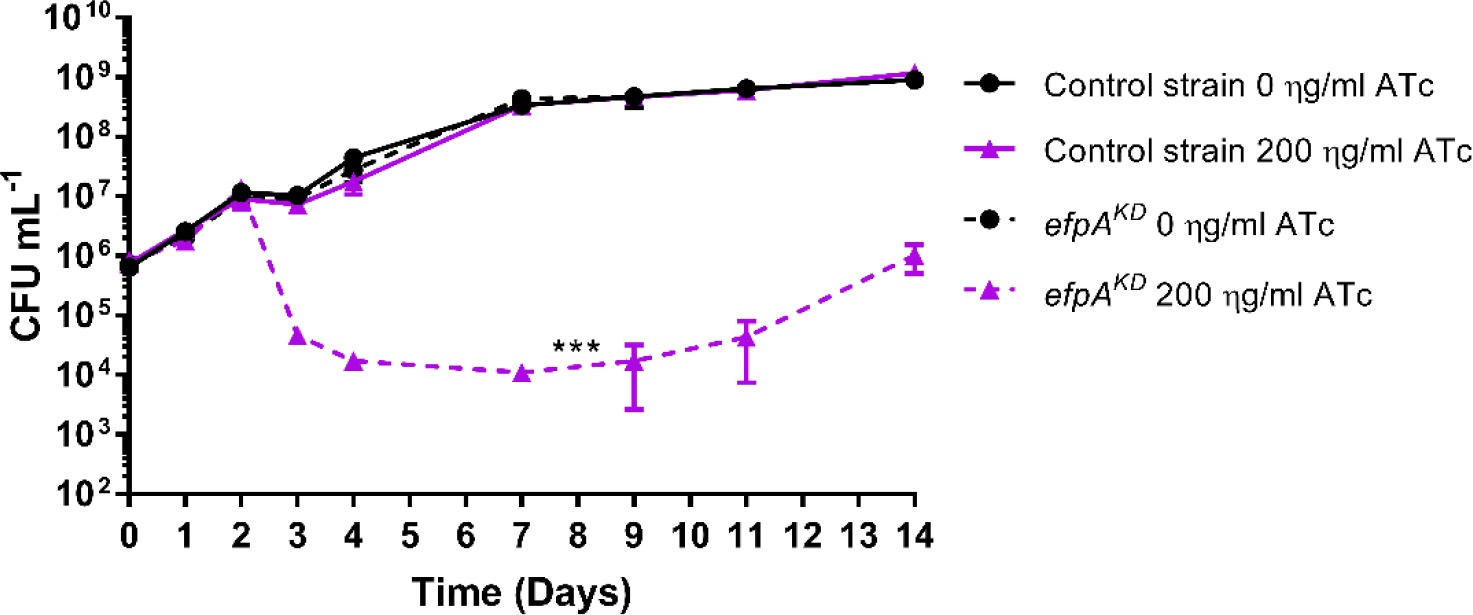
Comparison of viability (CFU mL^-1^) of the control strain (solid line) and *efpA*^KD^ (dotted line) over a 14-day time-course induced with (purple triangles) or without (black circles) 200 ng mL^-1^ ATc. The ATc was added to respective cultures once, at day 0. All values represent the mean of three biological replicates. Error bars are ± SD. Statistical significance was determined across the whole time-course by two-way ANOVA between each strain comparing the responses of each strain with and without ATc addition. *** = *P* ≤0.001. Non-significant differences are not displayed on graph. CFU mL^-1^ data was log_10_ transformed prior to statistical analysis.

### Induction of the CRISPRi system leads to a dramatic reduction in *efpA* expression and changes in cell morphology

To assess repression of *efpA* expression, the copy number of *efpA* mRNA transcripts was determined in *efpA*^KD^ with or without ATc-induction, using RT-qPCR. ATc was added at day 4 of log phase growth and *efpA* transcript copy numbers were quantified in comparison with non-induced cultures (Fig. 3). At day 1, *efpA* transcript levels were 43-fold lower in the induced *efpA*^KD^ cultures compared to the non-induced cultures, confirming the repression of *efpA* expression (Fig. 3). Following this, the transcript levels rose over time and were 3-fold lower in the induced *efpA*^KD^ by day 3 post-induction and recovered to equivalent expression levels in the non-induced cultures, by day 7.

**Figure 3:**
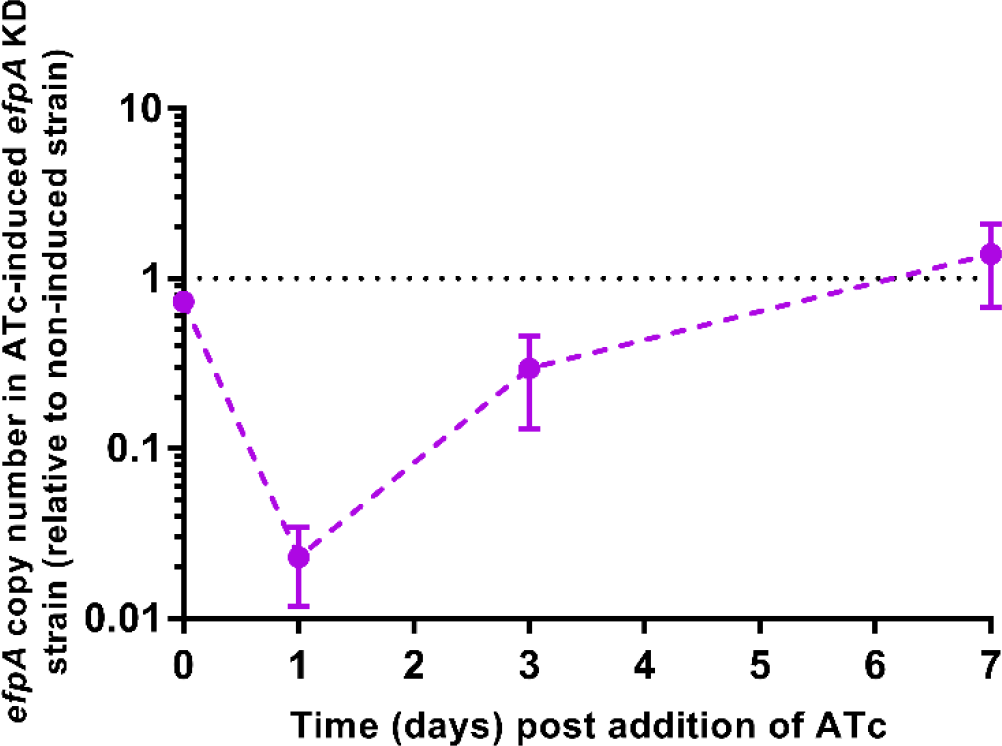
Repression of *efpA* in *M. tuberculosis* by CRISPRi, determined, by qRT-PCR of ATc-induced (200 ng mL^-1^) *M. tuberculosis efpA*^KD^ cultures relative to non-induced condition. ATc was added at day 4. Transcript copy numbers were determined by a qRT-PCR absolute quantification method. *efpA* copy number values were normalised relative to *sigA* copy number values across the time-course. Error bars are ±SD of the mean of three biological replicates.

To observe whether the repression of *efpA* had an impact on cellular morphology, scanning electron microscopy (SEM) was used to study the *efpA*^KD^ and the control strain, over time, in culture with or without ATc-addition at day 0. Images were taken on day 7 post ATc-induction. The SEM images showed no change in the control strain upon ATc-induction (Fig. 4A) and there were no morphological changes in *efpA*^KD^ cultures in the absence of ATc (Fig. 4B). However, when *efpA*, was repressed, an elongated phenotype was observed (Fig. 4C and 4D); many cells were extended up to 4-7 μm in length (Fig. 4C). Branched and blebbing morphology was also observed (Fig. 4D). These results, taken together, implied that there was an irregularity in the ability of the cells to divide when *efpA* expression diminished.

**Figure 4:**
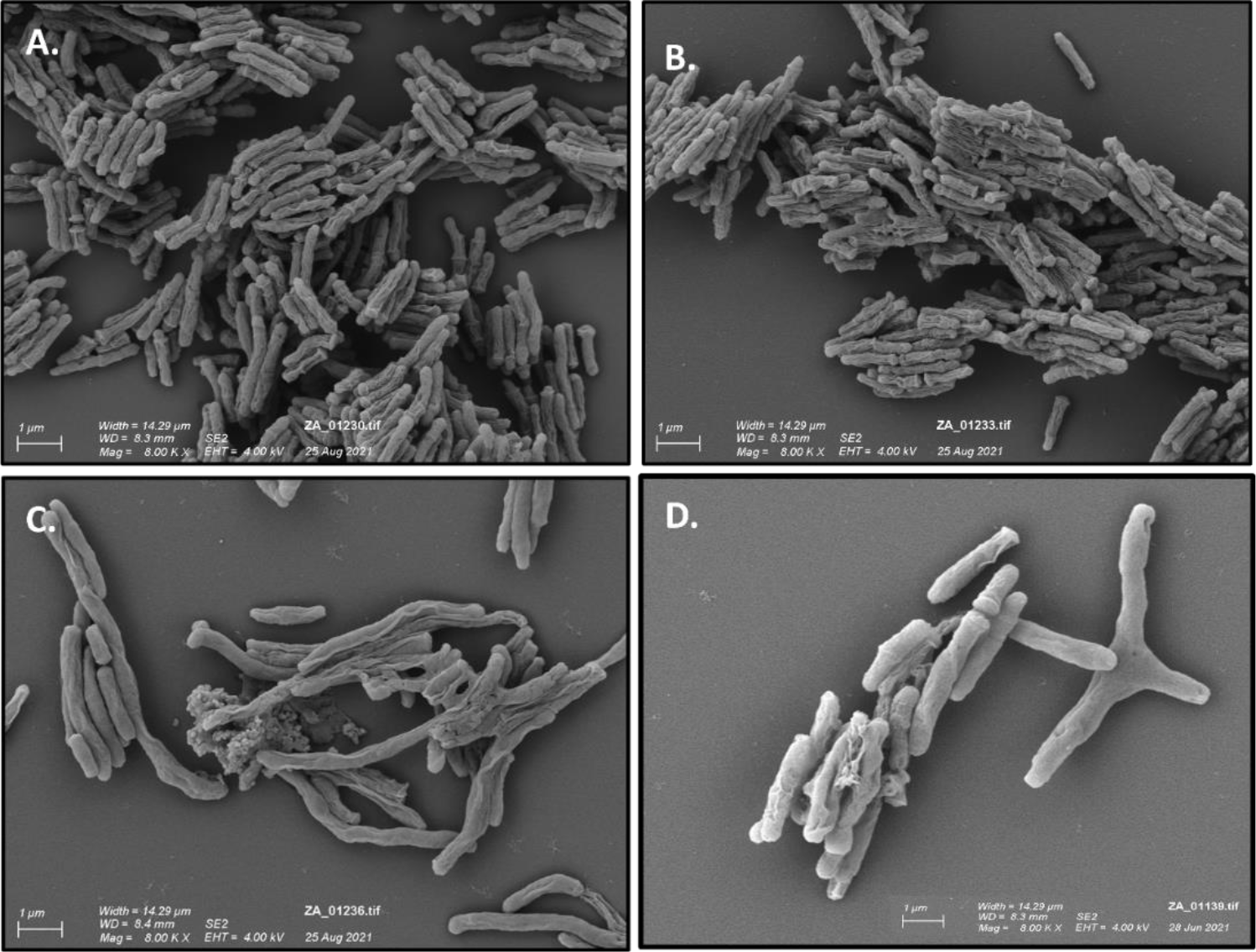
Scanning electron micrographs of control strain cultures exposed to 200 ng mL^-1^ ATc (A) and *efpA*^KD^ cultures in the absence of ATc (B) or exposed to 200 ng mL^-1^ ATc (C and D). Scanning electron micrographs were imaged at 8,000 x magnification.

### Repression of *efpA* prevents re-growth of *M. tuberculosis* during INH exposure

We determined whether *efpA* repression led to an increased sensitivity of *M. tuberculosis* to INH. The *efpA*^KD^ strain and the control strain were cultured for 45-days, in triplicate flask cultures, in Middlebrook 7H9 medium. At the point of inoculation, the following were added to cultures: either ATc at 200 ng mL^-1^, INH at 0.5 μg mL^-1^, a combination of INH/ATc, or no drugs at all. Samples were removed at days 0, 1, 2, 3, 4, 8, 10, 14, 17, 21, 24, 31, 38, 45, for total viable counts. The viable counts provided confirmation that the INH was imparting bactericidal effect and not the ATc, in these experiments. As observed in Fig. 2, in the absence of INH, the ATc-induced *efpA*^KD^ cultures, exhibited a reduction in viability due to *efpA*-repression (Fig. 5). In the presence of INH alone, both the *efpA*^KD^ and control strains exhibited an equivalent steep drop in viability (3logs_10_ CFU mL^-1^) over the first 3 days and started to recover by day 5, as expected from previously published *in vitro* INH-exposure experiments (de Steenwinkel *et al*., 2010; Jeeves *et al*., 2015; Hendon-Dunn *et al*., 2019). By day 24, both strains recovered to the same level of viability (CFU mL^-1^) that had been observed at the point of INH-addition. Cultures of the control strain were able to recover following the drop in viability after exposure to a combination of ATc and INH; following the same pattern of response as the control strain exposed to INH alone. For *efpA*^KD^, a combination of ATc and INH resulted in an initial decline in viability that was consistent with the response to INH alone. However, between days 7 and 14, *efpA*^KD^ cultures exhibited a severe reduction in viability and fluctuated at a level of viability around the limit of detection (10 CFU mL^-1^) over the remainder of the 45-day time course (Fig. 5). In conclusion, the ability of *M. tuberculosis* to recover from INH-exposure was abolished through the repression of *efpA*.

**Figure 5:**
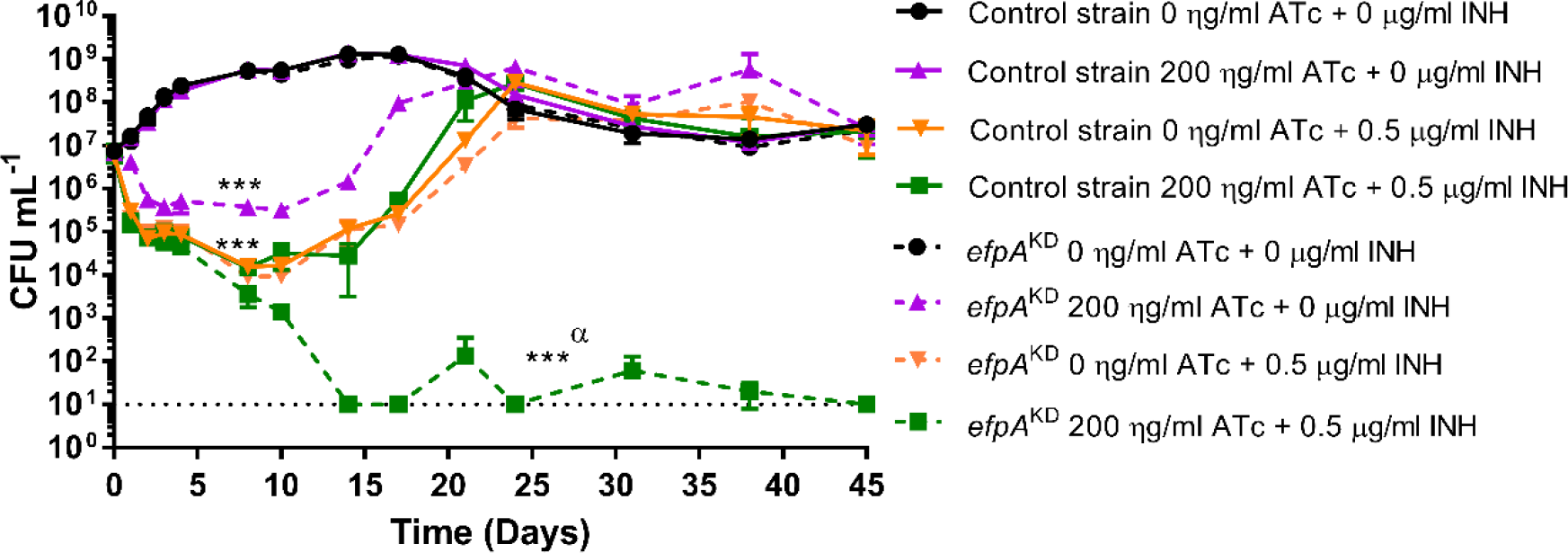
The viability of the control strain (solid lines) and *efpA*^KD^ (dotted lines) over a 45-day time-course, exposed to either 0.5 μg mL^-1^ of INH (orange inverted triangles), 200 ng mL^-1^ ATc (purple triangles), 0.5 μg mL^-1^ isoniazid and 200 ng mL^-1^ ATc in combination (dark green squares), or no drug (black circles). INH/ATc were added at culture inoculation (day 0). The *M. tuberculosis* response was measured by viable counts (CFU mL^-1^). All values represent the mean of three biological replicates. Error bars are ± SD. Statistical significance was determined by a three-way ANOVA between each drug condition and the drug-free condition/one drug condition in each strain. ***=*P* ≤0.001 and non-significant differences are not displayed on graph. ***α = a three-way ANOVA using CFU mL^-1^ values set at the limit of detection (10 CFU mL^-1^). CFU mL^-1^ data was log_10_ transformed prior to statistical analysis.

## Discussion

### Targeted repression of *efpA* using CRISPRi results in a loss of viability in *M. tuberculosis*

EfpA is a putative efflux pump that may be modulating the effectiveness of TB drugs. We used an inducible CRISPRi system to repress *efpA* expression in *M. tuberculosis* (Singh *et al*., 2016). This resulted in a statistically significant loss (P≤ 0.001) in the viability of *efpA*^KD^ in ATc-induced cultures compared to uninduced cultures and resulted in a triphasic response; bactericidal, followed by static, and finally re-growth, which could be explained by an initial loss of bacteria for which *efpA* is fully repressed, followed by a phase of non-replication, and finally regrowth as the effects of ATc-induction were waning (Fig. 2). This profile is reflected in the levels of *efpA* transcript overtime, being markedly reduced at 24h, with a restoration of transcript levels by day 7 post ATc-induction. Additional supporting evidence for non-replication was provided by the SEM images (Fig. 4), that revealed bacteria with increased cell length that appeared not to be dividing and exhibited branching or blebbing morphologies. Whole genome transcriptome analyses (data not shown) have highlighted the lower expression of genes that are required for cell wall biosynthesis and cell division, particularly at day 7, post-induction. Some examples being, *fipA, murD, murF, murX* and *ald*. These data taken together suggest a role for *efpA* in cell division/cell wall biosynthesis, which warrants further study.

### Repression of *efpA* prevented re-growth of *M. tuberculosis* during INH exposure

The focus of the work was to understand the role of *efpA* in the susceptibility of *M. tuberculosis* to antibiotics, the development of antibiotic resistance, and more specifically, recovery *in vitro*, when the organism is exposed to INH. Repression of *efpA* combined with exposure to INH caused a more pronounced reduction in viability compared to INH-alone and abolished the classic re-growth often observed for *M. tuberculosis* responses, *in vitro* (Fig. 5).

Re-growth after INH-exposure has been observed in a variety of settings including *in vitro* batch log-phase cultures, continuous culture conditions, murine and guinea pig studies, and in patients with pulmonary tuberculosis (Jindani *et al*.,1980; Klemens & Cynamon, 1996; Gumbo *et al*., 2007; Ahmad *et al*., 2009; de Steenwinkel *et al*., 2010; Jeeves *et al*., 2015; Hendon-Dunn *et al*., 2019). We have previously sought to understand the reasons for this re-growth, observed in continuous culture and addressed the hypothesis that re-growth was due to the persistence of drug-tolerant, slow growing bacilli. We found through a combination of two studies that the presence of isoniazid (singly and in combination with other frontline drugs) led to re-growth, irrespective of growth-rate or isoniazid concentration (Jeeves *et al*., 2015; Hendon-Dunn *et al*., 2019). Whole genome transcriptomics pointed to a variety of potential tolerance mechanisms, including upregulation of efflux pumps (Jeeves *et al*., 2015). In the current study, during the first few days of treatment, *efpA*-repression led to further reduction in the viable count (over INH-alone) but did not lead to an increased rate of killing rate by INH, but recovery of the culture was abolished. We observed previously in continuous culture that *efpA* was more highly expressed during the early bactericidal activity, at 2 days post-INH addition (Jeeves *et al*., 2015). Therefore, despite there being no additional impacts of *efpA*-repression on the initial rate of killing by INH, the organism may be preparing itself, by upregulation of *efpA*, during this phase, to enable survival, development of a tolerant and/or resistant population, for recovery later in its response to INH-exposure.

To interpret these findings, we need to firstly determine the physiological role of *efpA* in the absence of INH to understand the contribution of *efpA*-repression to the lack of re-growth during isoniazid exposure. Repression of *efpA* may lead to a reduction in the level of INH expelled from the cell resulting in an increased intracellular concentration, which in turn may hinder the ability of *M. tuberculosis* to develop INH-resistance mutations, as proposed by Schmalstieg *et al*. (2012). Further explanations could be that cell wall modifications observed due to *efpA*-repression may be enhancing the activity of INH. If we consider the mode of action of INH, an alternative explanation for increased INH activity could result from a shift in the NADH/NAD^+^ ratio. Activation of INH requires the formation of an INH-KatG-NAD+ adduct. It has been observed in *Mycobacterium tuberculosis* that altered NADH/NAD^**+**^ ratios lead to resistance to INH and ethionamide (Vilchèze *et al*., 2005). In *P. aeruginosa*, high intracellular levels of NADH contribute to efflux-mediated antibiotic resistance but can also drive cell death through an increase in reactive oxygen/nitrogen species (Arce-Rodríguez *et al*., 2022). We hypothesise that if EfpA was essential for maintaining the intracellular NADH/NAD^+^ ratio, when repressed, an imbalance in the ratio of NADH/NAD+ would lead to cell death through increased ROS. Further to these non-specific effects, an imbalance in the ratio of NADH/NAD+ could also alter the availability of NAD^+^ for isoniazid activation. The proposed hypothesis is speculative and warrants further investigation as there would be clear advantages of targeting EfpA through the combined effects of cell-mediated death and antibiotic activity. These investigations would include experiments to understand the interplay between EfpA-repression and the activity of other antibiotics.

### Is EfpA a good target for therapeutics?

This study provides further evidence that EfpA is essential in *M. tuberculosis* and may be a legitimate new target for drug discovery. The finding that repression of *efpA* prevents re-growth of *M. tuberculosis* after isoniazid exposure is significant, especially in retaining the use of isoniazid in current TB regimens. In the first instance, it would be useful to see how the inhibition of EfpA impacts pathogenesis and INH efficacy in animal models of *M. tuberculosis* infection and to see whether relapse is reduced. The early bactericidal activity (EBA) of INH reduces the bacterial burden substantially over the first two days during patient treatment. If EBA could be enhanced by an efflux breaker that improves INH efficacy further and removes re-growth/relapse, the lifespan of INH as a treatment of susceptible TB could be extended, whilst contributing to a reduction in the development of MDR, and the shortening of treatment times. EfpA was characterised in a recent chemical-genetic screening study Johnson *et al*. (2019 & 2020), which resulted in the development of two inhibitors that exhibited good inhibitory activity against EfpA, called BRD-8000.3 and BRD-9327. EfpA may also be involved in the efflux of Pretomanid, (a member of the nitroimidazole class of drugs), which is used in the BPaL combination for the treatment of MDR-TB), indicating that EfpA may be involved in the efflux of other drugs (Manjunatha, Boshoff, & Barry, 2009). Further studies will determine whether the inhibition of EfpA is a viable strategy to augment multi-drug therapy for tuberculosis by repressing efflux activities.

## Acknowledgments

Author contributions were as follows: The study was conceptualised by AR, SW, & JB; technical support for construction of the CRISPRi system was provided by VF and SK; technical support for the INH-exposure experiments was provided by CM & JB; experiments and data analyses were performed by AR; the manuscript was written by AR & JB; and reviewed/edited by all authors. This work was funded by the Department of Health, and Public Health England, UK. The views expressed in this publication are those of the authors and not necessarily those of the Department of Health or Public Health England. The authors acknowledge Howard Tolley and Katherine Davies for the scanning electron microscopy and thank Prof Andrew Gorringe for reviewing the manuscript prior to submission.

